# Environmental differences explain subtle yet detectable genetic structure in a widespread pollinator

**DOI:** 10.1101/2021.07.09.451741

**Authors:** Marcel Glück, Julia C. Geue, Henri. A. Thomassen

## Abstract

**Background:** The environment is a strong driver of genetic structure in many natural populations, yet often neglected in population genetic studies. This may be a particular problem in vagile species, where subtle structure cannot be explained by limitations to dispersal. These species might falsely be considered panmictic and hence potentially mismanaged. Here we analysed the genetic structure in an economically important and widespread pollinator, the buff-tailed bumble bee (*Bombus terrestris*), which is considered to be quasi-panmictic at mainland continental scales. We first quantified population structure in Romania and Bulgaria with spatially implicit Fst and Bayesian clustering analyses. We then incorporated environmental information to infer the influence of the permeability of the habitat matrix between populations (resistance distances) as well as environmental differences among sites in explaining population divergence.

**Results:** Genetic structure of the buff-tailed bumble bee was subtle and not detected by Bayesian clustering. As expected, geographic distance and habitat permeability were not informative in explaining the spatial pattern of genetic divergence. Yet, environmental variables related to temperature, vegetation and topography were highly informative, explaining between 33 and 39% of the genetic variation observed.

**Conclusions:** Where in the past spatially implicit approaches had repeatedly failed, incorporating environmental data proved to be highly beneficial in detecting and unravelling the drivers of genetic structure in this vagile and opportunistic species. Indeed, structure followed a pattern of isolation by environment, where the establishment of dispersers is limited by environmental differences among populations, resulting in the disruption of genetic connectivity and the divergence of populations through genetic drift and divergent natural selection. With this work, we highlight the potential of incorporating environmental differences among population locations to complement the more traditional approach of isolation by geographic distance, in order to obtain a holistic understanding of the processes driving structure in natural populations.

## Background

Detecting genetic population structure and its underlying causes is crucial to better understand basic evolutionary ecological processes and how these are affected by human actions, as well as to improve conservation strategies. Population structure is often inferred using methods that are only based on genetic data, and do not take into account the geographic relationships between populations (1). These methods perform well when population structure is strong, but may fail to correctly detect weak structure. Considering spatial relationships can help to improve the detectability of weak structure (2–4), but mainly when there are small but distinct genetic breaks in geographic space. Gradually changing genetic population structure is, however, notoriously difficult to detect, in particular when structure is subtle. As a consequence, species exhibiting such patterns might incorrectly be considered panmictic.

Population structure is the result of a balance between gene flow, genetic drift, natural selection, and mutations. As the likelihood of successful dispersal between populations decreases with geographic distance, gene flow starts to become limited and populations diverge through genetic drift. Under this scenario, a positive relationship between geographic and genetic distance is anticipated, a pattern termed isolation by distance (IBD, 5). Moreover, heterogeneous conditions of the habitat through which dispersal takes place impose varying levels of resistance to dispersing individuals. They may therefore not follow a straight line, but instead the path of least resistance. Least-cost path (6) and isolation by resistance (IBR, 7) analyses aim to quantify this heterogeneity in habitat permeability and its effect on population structure. Both IBD and IBR describe processes resulting in neutral population divergence, and are jointly coined isolation by dispersal limitation (8). In addition, species experience heterogeneous environmental conditions that may exert strong selection pressures on populations, potentially leading to local adaptation and hence population divergence (9, 10). Interestingly, the prolonged reduced fitness of dispersers (11) that are maladapted to newly encountered conditions might result in the disruption of genetic connectivity. As a consequence, populations may differentiate by means of genetic drift in neutral markers, a pattern termed isolation by environment/ecology (IBE, 12, 13) or, alternatively, isolation by adaptation (IBA, 14). Approaches that neglect environmental heterogeneity as a driver of population structure may thus be overly simplistic and result in an incomplete picture of the processes that structure natural populations (e.g. 15, 16), failing to recognize subtle and gradual patterns of genetic turnover.

To this end, the field of landscape genetics (17–19) provides the tools and data to not only analyse genetic information in a spatially explicit context, but to also consider local environmental conditions and those of the habitat matrix between populations in explaining non-random gene flow across the landscape (11). Thus, it has become possible to study the full range of evolutionary ecological processes driving population divergence, to tease apart their relative importance, and to take advantage of the availability of spatially continuously varying environmental data to detect subtle population structure.

Integrating environmental dissimilarities into the analysis of population structure is particularly promising in species capable of dispersing widely. Typically, these species do not show genetic patterns consistent with IBD or IBR, and exhibit divergence levels close to panmixia (20, 21). Such a species, which is deemed quasi-panmictic at the subspecies level, is the buff-tailed bumble bee (*Bombus terrestris*). This highly polymorphic pollinator occurs across various environmental gradients (22, 23), and morphological differences have prompted a division in nine subspecies (24). Studies on the population structure in this species have inferred little to no genetic divergence at spatial scales ranging from small-scale assemblages (25) over countries (26) to continents (27, 28). Exceptions are populations separated by strong oceanic barriers on islands or between continents (27–30) where pronounced structure can be observed. Although large water bodies and strong winds have been implied to limit gene flow (28), the potential for subtle population structure and how it might be influenced by environmental heterogeneity remain unelucidated.

Here we aimed to quantify population structure in the buff-tailed bumble bee in two countries that exhibit pronounced landscape heterogeneity that is readily exploited by this species. We hypothesized that this species shows genetic structuring consistent with a scenario of IBE. Further, as the buff-tailed bumble bee is highly vagile (23, 31), we expected geographic distance and landscape resistance to play only minor roles in explaining population divergence.

We used both spatially implicit and explicit analyses to identify the most likely drivers of population structure. Deploying 12 highly polymorphic microsatellite loci, we did not detect population structure using spatially implicit Bayesian clustering. However, Fst analyses demonstrated subtle yet significant genetic divergence. Using generalized dissimilarity modelling (GDM, 32), simultaneous analysis of the influence of geographic distance, landscape resistance, and environmental dissimilarities among populations demonstrated subtle and negligible influence of geographic and resistance distances on population structure. Yet environmental dissimilarities were highly informative in detecting and unravelling the drivers of genetic structure in this widespread and abundant pollinator.

## Results

### Genotyping and exclusions

A total of 376 out of 385 buff-tailed bumble bees were successfully genotyped at 12 microsatellite loci (Additional file 1: Table S1), with locus-specific error rates ranging between 0 and 5.56%. Using GenAlEx and Colony, we detected and excluded 8 clones, 19 full siblings, and 2 individuals inferred as being clones and full siblings at the same time. Following the identification of males based on multilocus heterozygosity scores, we excluded 36 putative males of which 34 were homozygous across all 12 loci and 2 across 10 or 11 loci.

### Descriptive statistics

Using Micro-Checker, we detected null alleles at all loci except for two loci (ms66 and ms86; Additional file 1: Table S1) with numbers ranging from 1 to 10. Signals of stuttering were present at five loci (ms39, 80, 81, 85, and 41) in one to eight populations. We did not detect signals of large allele dropout. Deviations from Hardy-Weinberg equilibrium (HWE) were observed in Genepop on the Web, with 8 out of 12 loci showing significant departure from HWE in 1 to 5 populations. We inferred overall significant (*P* = 0.027) linkage disequilibrium (LD) for the locus pair ms39–ms80, which was, however, not supported by population pairwise analyses, where significant LD was detected only in the populations Drӑgusani and Valea Hotarului. As null alleles, stuttering, and deviations from both HWE and LD were not consistent across either loci or populations, we retained all loci in the data set.

Genetic diversity exhibited by the bumble bee populations was medium to high, with observed heterozygosity (HO) ranging from 0.46 to 0.60 in a data set containing diploid females only (diploid data set; ‘dpds’) (Additional file 1: Table S2). After correcting for unequal sample sizes by rarefaction for 7 individuals, the number of alleles ranged from 2.70 to 3.09 for ‘dpds’ and from 2.26 to 3.08 for a data set comprising haploid males and diploid females (mixed-ploidy data set; ‘mpds’). Overall population divergence was low in both data sets (global Fst = 0.006 (*P* = 0.02) and 0.041 (upper 95% CI: 0.047) for ‘dpds’ and ‘mpds’, respectively). In population pairwise comparisons, Fst values ranged from 0 to 0.07 (‘dpds’, Additional file 1: Table S3) and from 0.01 to 0.12 (‘mpds’, Additional file 1: Table S4). Following FDR correction of *P* values for ‘dpds’, five pairwise comparisons remained borderline significant (*P* = 0.053–0.059) with Fst values between 0.046 and 0.065. None of the ‘mpds’ Fst values surpassed its corresponding upper 95% confidence interval.

### Genetic clustering

Based on visual interpretation of the obtained bar plots, Structure runs using the Admixture and Correlated Allele Frequency models with population IDs as priors, suggested the absence of clear genetic structure (*K* = 1). This pattern was consistently inferred in all five independent runs for each *K*, ranging from 2 to 25 (Additional file 2: Figure S1). Excluding populations with fewer than 10 individuals (‘pop10’) did not change this conclusion, regardless of whether individuals were initialized from their respective populations or not (results available on Dryad).

Despite the potential occurrence of two subspecies (*B. t. dalmatinus* and *B. t. terrestris)* native to Romania and Bulgaria (33), it seems unlikely that *B. t. terrestris* was included in this analysis, as its known range is limited to the western border regions. Additionally, we would expect its inclusion to result in pronounced clustering, something we did not observe here. In summary, using Bayesian clustering, the buff-tailed bumble bee resembles a large panmictic population across Romania and Bulgaria, even when sampling locations were used as a prior for potential structure.

### Landscape genetic analyses

To further investigate whether subtle population structure could be detected when habitat characteristics were included in our analyses, and to estimate the relative effects of the potential drivers of structure, we ran generalized dissimilarity models (GDMs, 32). Local habitat conditions were characterized using a set of topographical, climate and vegetation variables, and resistance distances were based on a species distribution model (SDM) generated previously (22). GDMs explained ~ 33.5 and 39.2% of the divergence observed for ‘dpds’ and ‘mpds’, respectively (Table 1). We compared these values to those of 1,000 random models. Although the random models performed surprisingly well and explained ~ 24.3 (lower CI–upper CI: 23.8–24.8) and 31.8% (31.2–32.3) of the divergence for ‘dpds’ and ‘mpds’, respectively, they were outperformed by the full models. Environmental dissimilarities contributed most to explaining the genetic variation. In contrast, geographic distance and the SDM-derived resistance distances performed poorly; they were not retained in the full models and only explained 0.00–0.07 and 0.00–0.01% of the total variation when analysed in isolation.

**Table 1:** Percent variance explained by the generalized dissimilarity models (GDMs) for the data set encompassing diploid (‘dpds’), and both haploid and diploid (‘mpds’) individuals.

|  | dpds | mpds |
| --- | --- | --- |
| Full | 33.48 | 39.18 |
| Env only | 33.48 | 39.18 |
| Geo only | 0.07 | 0.00 |
| Res only | 0.00 | 0.01 |
| Random | 24.30 | 31.78 |
| Lower CI | 23.77 | 31.24 |
| Upper CI | 24.82 | 32.32 |

Models were run with five different input data sets. (1) Full: geographic distance, resistance distance and environmental variables; (2) Env only, (3) Geo only, (4) Res only: environmental variables, geographic distance, and resistance distance only, respectively. (5) Random: mean of 1,000 models with randomly generated environmental variables. Lower/Upper CI: lower/upper 95% confidence interval of the random models.

The environmental variables retained included temperature, precipitation, topography, measures of surface moisture, and vegetation density (Additional file 2: Figures S2, S3). The most important variables in explaining genetic turnover in both ‘dpds’ and ‘mpds’ were slope, Leaf Area Index (LAI; mean and standard deviation) and mean temperature of the coldest quarter (Bio 11). Seasonality in surface moisture (QSCAT (seasonality)), isothermality (Bio 3) and mean temperature of the wettest quarter (Bio 8) contributed as well, but their importance varied between the two data sets. Predicted genetic turnover across Romania and Bulgaria was most pronounced along elevation gradients, such as between the Danube Delta and the Carpathians in Romania and the Balkan Mountains in Bulgaria (Figure 1 (b)).

**Figure 1:**
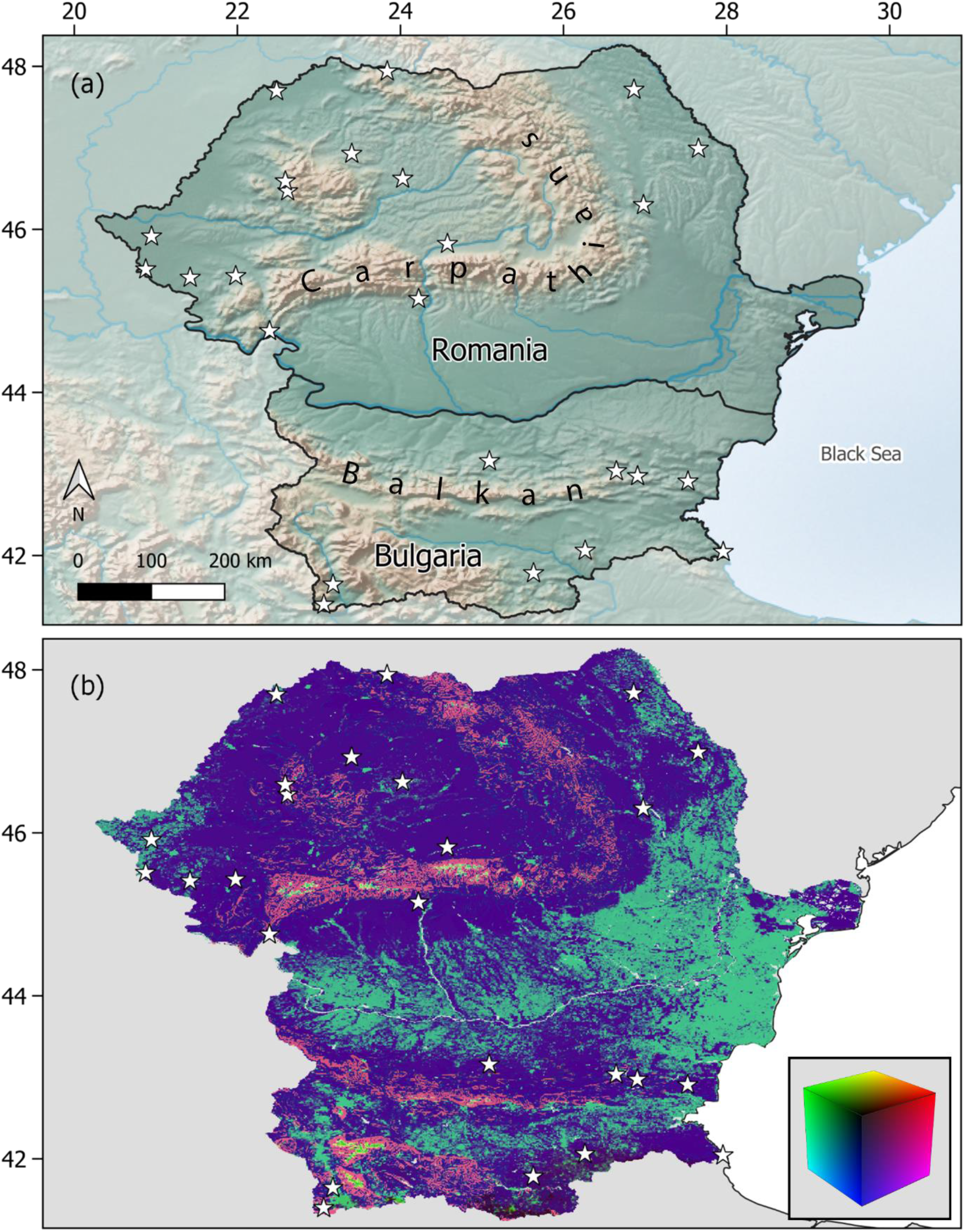
Study region and spatial GDM prediction. (a) Location of the study area in South-Eastern Europe. Made with Natural Earth (130). (b) Prediction of genetic turnover across Romania and Bulgaria, where increasingly dissimilar colours indicate more pronounced turnover. Stars mark population locations.

## Discussion

Incorporating environmental data in population genetic studies might help to identify and explain subtle population structure in vagile species, whose dispersal is usually not constrained by geographic distance or the habitat through which dispersal takes place. To assess the informativeness of environmental data, we quantified population structure in the widespread and abundant buff-tailed bumble bee, a species that shows subtle to no structure at mainland continental scales. Using genetic data only, we inferred near-panmixia across two Eastern European countries, Romania and Bulgaria, showing pronounced landscape heterogeneity. With a landscape genetics approach, we related genetic divergence to turnover in environmental variables. Geographic distance was not informative, nor were resistance distances derived from a species distribution model (SDM), that quantifies resistance to dispersal. Population structure in buff-tailed bumble bees thus follows a pattern of isolation by environment, where dissimilar habitats reduce the fitness of dispersing individuals, resulting in a disruption of genetic connectivity to a level that it cannot completely balance the effects of genetic drift.

The subtle level of divergence observed among Romanian and Bulgarian populations agrees with a previous study demonstrating weak but significant genetic structure among buff-tailed bumble bee populations in continental Europe (28). Although some of the genetic divergence observed in the aforementioned study might simply be attributed to the inclusion of three subspecies (33), together these studies highlight the possibility of weak but detectable genetic structure after previous work had repeatedly concluded that the buff-tailed bumble bee exhibits insignificant levels of genetic differentiation and hence population panmixia (25–27). In general, overall low population structure has not only been detected in *B. terrestris*, but seems fairly common among *Bombus* species, including but not limited to *B. lapidarius* (34), *B. hortorum*, *B. ruderarius*, *B. soroeensis* (35) and *B. ignitus* (36). Quasi-panmixia in these species is likely driven by extensive gene flow (23, 27, 31), strong enough to override most of the divergence caused by genetic drift and divergent natural selection. This hypothesis is in line with the absence of detectable levels of isolation by distance in our study as well as mainland populations of many other *Bombus* species, such as *B. pascuorum* (37), *B. vosnesenskii* (38), *B. lapidarius*, *B. hortorum*, *B. ruderatus (39)* and *B. flavifrons* (40). Dispersal, and hence gene flow, may not follow a straight line, but rather a path of least resistance through suitable habitat. Resistance distances are thus often considered to be a better proxy of between-population dispersal than geographic distance. Nevertheless, our analyses suggest that gene flow is not constrained by variation in habitat permeability either, a finding potentially caused by the absence of extensive oceanic barriers in the study area (41–43).

Subtle genetic structure might also result from the buff-tailed bumble bee‘s generalist foraging behaviour and the production of many workers (44–46), both potentially allowing this species to efficiently exploit natural and semi-natural habitats unsuitable for related species with a narrower dietary niche (47, 48). In this way, the buff-tailed bumble bee’s broad niche might translate to a more continuous distribution of nests across the landscape, with ample opportunity for gene flow among populations. However, with increasing nest densities, the benefit of a more continuous distribution might increasingly be cancelled out by strong density-induced intraspecific competition for forage (49, 50), which may reduce overall nest performance and thus reproductive output (51, 52). Hence, compared to related species, the lower nest densities observed in *B. terrestris* (39, 53, 54) might even benefit gene flow, as a relaxation of competition might allow more nests to contribute to genetic connectivity, resulting in an overall weak structure in this species.

Although quite subtle, the divergence we observed was best explained by environmental dissimilarities, in particular through the mean temperature of the coldest quarter, Leaf Area Index (LAI) and slope. The importance of temperature did not come as a surprise, as it is an important predictor of the distribution of *Bombus* species (22, 55) and known to govern the emergence time of queens from hibernation (56). Hence, queens hibernating in warmer areas might emerge earlier than those in colder areas. In fact, asynchronous emergence times may translate to phenological mismatches and hence reproductive isolation between early and late colonies. Even though laboratory experiments with *B. perplexus* and *B. lucorum* did not support this hypothesis (57, 58), the complex environment these species experience throughout their life cycles is unlikely to be fully reproduced in the laboratory (59). Indeed, emergence patterns of sexuals in natural populations of several species, including *B. flavifrons* and *B. lucorum/terrestris*, differed strongly (60, 61), highlighting the plausibility of this hypothesis. As buff-tailed bumble bee gynes usually only mate once (62, 63), the asynchronous emergence of sexuals allows early emerging males to effectively monopolize queens, thus promoting population divergence.

Even though differences in local temperatures might thus facilitate genetic divergence, the influence of this environmental variable on the species’ life cycle is more nuanced, potentially even allowing the variable to contribute to genetic homogeneity. Indeed, facultative endothermy (64) and the ability for collective thermoregulation (65) allows nests to survive even under low temperatures (66), enabling great phenological flexibility (67) and a change from univoltism to bivoltism in regions hitherto considered incompatible with this phenological pattern (52). Notably, this change in phenology seems to spread northwards into colder regions (52), a pattern coinciding with northward range shifts in this species (68). As a bi-annual iteration through the species’ life cycle effectively doubles the number of potential admixture events, it might contribute to genetic homogeneity. However, generally cold winters in Romania and Bulgaria (69, 70), and the absence of sufficient forage during the cold season might limit bivoltism to a few inland and coastal regions in both countries (70).

Even though mean LAI was retained as an important variable in the final models, the mechanism through which this measure of greenness is associated with genetic structure remains unclear. Yet, mean LAI was highly correlated with percent tree cover (r = 0.83, Additional file 1: Table S6), and although forests do not seem to limit bumble bee movement (71, 72), buff-tailed bumble bee queens prefer open habitat for nesting (73). Forests may thus reduce genetic connectivity by constraining the amount of available nesting habitat (41, 74). Moreover, assuming a negative effect of woodland on the range expansion of this species allowed to accurately model its invasion pattern in Japan (75), suggesting that queen-borne range expansion might indeed be limited by forests.

The slope of the terrain, inferred here as influential in shaping genetic divergence, might be an important determinant of suitable hibernation habitat (56, 76). However, the distribution of bumble bee populations across the landscape reflects the distribution of suitable nesting habitat, and only coincides with hibernation locations if queen dispersal after hibernation is strongly limited (77). Given that buff-tailed bumble bee queens are instead highly vagile (23, 78), the mechanism of how slope governs genetic divergence remains unclear.

Although we included a large set of environmental variables, about 60% of the genetic variation remained unexplained. As we aimed to unravel the factors shaping and maintaining large-scale genetic structure in this species, divergence explained by small-scale processes, temporal fluctuations or colony-intrinsic traits were beyond the scope of this work. Genetic divergence might also result from demographic processes, such as bottlenecks (36), which we could not cover here. Future studies might also investigate the influence of the abundance and the spatial arrangement of plant species producing nectar or pollen in high quantities (59, 79, 80) on structuring bumble bee populations. Finally, habitat alterations such as intensified farming practices might also structure populations, in particular through a synergy with natural stressors (81, 82).

## Conclusions

Seemingly panmictic populations might exhibit subtle genetic structure that can only be detected and understood when considering the environment as a potential driver of structure. We inferred subtle divergence in a widespread and abundant, quasi-panmictic pollinator that was not explained by geographic distance or variation in habitat permeability. Yet, using a suite of environmental variables, we showed for the first time in this species that environmental dissimilarities are informative in explaining spatial patterns of genetic structure. We encourage scientists to consider environmental data in their analyses to detect genetic structure and uncover its underlying drivers.

## Methods

### Study species and study area

The buff-tailed bumble bee (*Bombus terrestris*) is a widespread and abundant pollinator species. Its native distribution covers much of the Palearctic realm, including Europe, North Africa, and the British and most Mediterranean and Atlantic islands (27, 30). Even within regions, the species exhibits a remarkable niche breadth and is present across a wide range of habitats (22, 83). Its polylectic foraging and high pollination efficiency (24, 84) has rendered this species a prime candidate for pollination in the greenhouse industry where it is deployed in high numbers (85). Upon introduction outside its native range, this species has spread rapidly and established itself in many countries including Chile, Japan and New Zealand (23, 84, 86–88), which highlights its potential to rapidly adapt to novel environments.

We conducted this study in Romania and Bulgaria, neighbouring countries in South-Eastern Europe that exhibit high heterogeneity in topography, climate, and land use. On a large scale, extensive mountainous areas with peaks up to 2,500 m shape the face of both countries. In Romania, the Carpathian Mountains predominate (89), whereas in Bulgaria the landscape is structured by alternating bands of high and low terrain that stretch from east to west across the country (90). The topography gives rise to various climatic zones ranging from alpine and subarctic to humid subtropical (89, 91). The landscape is a mosaic of natural areas such as plains, open woodland, and extensive forests, as well as inhabited and in- and extensively used agricultural areas. This pronounced fine- and large-scale spatial turnover provides an ideal study ground to identify subtle population structure and its drivers in a species believed to be quasi-panmictic.

### Field sampling

Over 5 consecutive years, from 2013 to 2017, we obtained 385 individuals from 25 locations across Romania and Bulgaria (Figure 1 (a)). Sampling sites (Additional file 1: Table S7) were at least 20 km apart to avoid overlapping foraging ranges (71, 92, 93). Locations spanned a wide range of habitat conditions, encompassing both natural and semi-natural habitats, as well as extensive environmental gradients with respect to climate, vegetation and altitude. Sites were visited only once by a small team of 2–3 people for approximately 1.5 hours each. Individuals regardless of sex were caught using insect nets and sacrificed in a jar with ethyl acetate (94). Subsequently, specimens were transferred to individual tubes containing 96% ethanol and stored at −20 °C after returning to Tübingen University.

### Marker choice, DNA extraction, and genotyping

Despite the increased application of high-throughput sequencing approaches, microsatellites remain a powerful yet cost-efficient way of detecting even subtle population structure. Interestingly, following disruption of genetic connectivity due to reduced fitness of dispersers, drift-induced divergence patterns, related to local adaptations, are expected to appear even in neutral markers as a pattern of IBE/IBA (12–14). Thus, microsatellites can be used indirectly to study the influence of the environment on shaping population structure in a widespread species such as *B. terrestris*.

From each individual, we extracted DNA from one or two legs using the DNeasy Blood and Tissue Kit and QIAamp DNA Micro Kit (Qiagen, Hilden, Germany). We followed the manufacturer’s protocols except for adding 20 μl of 1 M dithiothreitol (DTT) solution to each sample, which facilitates the extraction of DNA from chitinous materials (95). Individuals of *B. terrestris,* specifically workers (96), can be difficult to distinguish morphologically in the field from another closely related bumble bee species, the white-tailed bumble bee (*Bombus lucorum)* (97). We therefore confirmed species identity using a 1,043 bp long fragment of the cytochrome c oxidase subunit I (CO1) gene (22).

We then amplified 12 previously developed (98) microsatellite loci in 3 multiplex reactions (PM1–PM3, Additional file 1: Table S1). PCR amplification was run in a total volume of 10 μl consisting of 5 μl PCR master mix (Qiagen), 2.1 μl HPLC H2O, 0.4 μl BSA (bovine serum albumin, 10 mg/ml), 1 μl primer solution (100 μM, Applied Biosystems) and 1.5 μl sample DNA. Samples were initially denatured at 95 °C for 15 min, followed by 25 cycles of denaturation (94 °C, 30 s), annealing (PM1: 56 °C, PM2/3: 60 °C, both for 90 s) and extension (72 °C, 60 s). An additional 20 cycles were run using the following settings: denaturation (94 °C, 30 s), annealing (44 °C, 90 s) and extension (72 °C, 60 s). PCR products were visualised on an ABI3730XL capillary DNA sequencer (Applied Biosystems) using a GeneScan 500 LIZ size standard (Applied Biosystems) at Macrogen Europe (The Netherlands). Results were analysed using GeneMarker v.2.4.0 (SoftGenetics, State College, PA). Samples that had not amplified successfully or for which scoring had not yielded conclusive results were re-amplified and re-scored. Individuals that repeatedly failed to amplify or yielded inconclusive results for the second time were excluded. The presence of genotyping errors was assessed by re-amplifying and re-scoring 36 randomly selected samples, representing approximately 9.5% of all individuals. Scoring results were compared between the first and second run and the mean error rate for each locus was calculated in Microsoft Excel.

### Data analysis

#### Identification and removal of clones, sibling workers and drones

As the presence of clones and full siblings is known to distort estimates of population structure, we first identified clones using the ‘Multilocus Matches’ analysis in GenAlEx v.6.503 (99, 100). Second, full siblings and additional clones were detected using the maximum likelihood approach implemented in Colony v.2.0.6.5 (101), which has been deemed the most reliable method for assigning sibship in bumble bees (102). In brief, we assumed a polygamous mating system to allow Colony to infer the relationship among all individuals entered as offspring, as well as inbreeding (103), the presence of clones, and dioecious reproduction with haplodiploidy. Two runs, differing in their seed values, were conducted with medium length using the full likelihood method with medium precision. Dropout rate was set to 0.001 and locus-specific mean error rates ranged from 0 to 5.56%. Individuals were considered clones or full siblings when they were inferred in both runs with probabilities > 0.8 and originated from the same population. For each inferred clone or full sibling pair, one randomly selected individual was retained, resulting in the so-called ‘complete data set’ (‘cpds’, Additional file 1: Table S7). As subtle genetic distances might not be inferred accurately for small populations (104), we excluded the populations ‘Billed’ and ‘Coastra’, resulting in a ‘mixed-ploidy data set’ (‘mpds’, Additional file 1: Table S7). Additionally, because haplodiploidy in *Bombus* deflates measures of heterozygosity, we identified and excluded putative males in ‘mpds’ based on observed multilocus genotypes. More specifically, individuals were considered males (i.e. haploid) if they were heterozygous for a maximum of 2 loci, resulting in a data set encompassing diploid individuals only (‘dpds’, Additional file 1: Table S7) from 21 populations throughout Romania and Bulgaria.

#### Population genetic analyses

After excluding putative clones, full siblings and haploid males, we tested for null alleles, stuttering and large allele dropout in Micro-Checker v.2.2.3 (105), applying 3,000 randomisations and a Bonferroni correction while omitting missing data. Hardy-Weinberg equilibrium (HWE) and genotypic linkage disequilibrium (LD) were assessed through Genepop on the Web v.4.2 (106) using the Marcov chain method with 10,000 dememorisations, 1,000 batches and 10,000 iterations per batch. Subsequently, we used GenAlEx v.6.503 (‘dpds’) and SPAGeDi v.1.5 (107) (‘mpds’) to compute genetic diversity indices, including observed and unbiased expected heterozygosity. Rarefied allelic richness (108) was obtained in hp-rare v.1.1 (109) (‘dpds’) and SPAGeDi (‘mpds’). We then computed population pairwise and global genetic distances (Fst) and corresponding *P* values for ‘dpds’ using GenAlEx with 9,999 permutations. Negative Fst values were converted to zero and *P* values were adjusted for multiple testing through false discovery rate (FDR) correction (110) using the ‘p.adjust’ function in R v.3.6.0 (111). For ‘mpds’ both the Fst matrix and global Fst value were computed with the ‘calcPopDiff’ function based on allele frequencies calculated with the ‘simpleFreq’ function in the polysat v.1.7.4 R package (112). Additionally, for both global and pairwise values, we computed the 95% confidence intervals of 10,000 bootstrap replicates. Fst values were considered significant if they surpassed the upper 95% confidence interval.

Following the computation of Fst estimates, we assessed genetic structure in ‘cpds’ through Bayesian clustering in Structure v.2.3.4 (1) using the Admixture and Correlated Allele Frequency models with population IDs as priors (113). We set the number of clusters (*K*) from 2 to 25 (number of populations sampled) and computed 5 iterations with 100,000 burnin and 1,000,000 Marcov chain Monte Carlo (MCMC) repetitions. We set INFERALPHA to 1, prompting Structure to infer the relative admixture between populations. Additionally, as uneven sampling sizes among populations might results in an incorrect number of clusters (114), 2 further Structure analyses were run after excluding populations with fewer than 10 individuals (‘pop10’, Additional file 1: Table S7). After setting the maximum *K* to 23, the first run was computed with the settings specified above while for the second run we additionally set STARTATPOPINFO to 1 to initialize each individual to its own population (115). Structure output was visualized with the pophelper v.2.3.1 R package (116).

#### Environmental variables and permeability of the habitat matrix

We characterized environmental conditions across Romania and Bulgaria using a set of 16 environmental variables (22) (Additional file 1: Table S5). This set included variables related to climate, vegetation and topography, after highly correlated variables had been excluded through variance inflation factor (VIF) analysis. To obtain a measure of habitat permeability, we further calculated population pairwise resistance distances based on the species distribution model (SDM) of *B. terrestris* (22). Computations were carried out in Circuitscape v.4.0.5 (117) with the SDM surface being treated as a conductance map, a cell size of 0.0083333 ° (i.e. 30 arcsec) and a cell connection scheme of 8 neighbours. Resulting population pairwise resistance distances (Additional file 1: Table S8) were used as an additional predictor in subsequent analyses.

#### Landscape genetic analyses

We established associations between genetic data and environmental variables using generalized dissimilarity modelling (GDM, 32) implemented in the gdm R package v.1.3.11 (118). The usefulness of this approach in landscape genetic analyses is widely acknowledged, as it allows fitting non-linear relationships between predictor and response variables. Additionally, the effects of geographic distance and the environment on genetic divergence can be analysed simultaneously, and estimates of variable importance are provided through permutation (119–121). Predictors inferred as informative in explaining genetic turnover yield I-splines that provide two pieces of information. The maximum height of the curve indicates the amount of biological change along the gradient, while the spline‘s shape informs about the rate of genetic turnover (120). In this study, we used this framework to infer the relationship between pairwise genetic distances (Fst, scaled between zero and one) and environmental variables, geographic distance and resistance distance. In total, we computed five different models, that included (1) environmental variables, geographic and resistance distances (full model), (2) environmental variables only, (3) straight-line geographic distances only, (4) resistance distances only, and (5) 1,000 models each using 16 random environmental variables to evaluate the performance of the full model. We considered the full model significant if its variation explained surpassed the 95% confidence interval of the random models. Subsequently, following the approach by Fitzpatrick and Keller (120), we used the inferred relationships between predictor and response variables at sampling sites to predict genetic turnover across Romania and Bulgaria. First, for each retained environmental variable, we extracted its value across the study area at 30 arcsec resolution. Using the ‘gdm.transform’ function from the gdm package, we then transformed environmental variables into ‘genetic’ importance values. We selected the three most influential and uncorrelated factors through principal component analyses (PCA) using the ‘princomp’ function with calculations performed on the covariance matrix. We purposely centred, but not scale-transformed the PCA to preserve differences in the magnitude of the genetic importance among environmental variables. With the ‘rasterize’ function in QGIS v.3.16.2 (122) we converted the obtained point values to rasters, merged them with the ‘r.composite’ function to a composite and assigned the corresponding PC scores to the RGB palette. The result visualizes differences in genetic composition, where increasingly dissimilar colours represent higher predicted genetic turnover.

## List of abbreviations

A_R_: Allelic richness
arcsec: Arcsecond
Bio 3: Isothermality
Bio 8: Mean temperature of the wettest quarter
Bio 11: Mean temperature of the coldest quarter
bp: Base pair
BSA: Bovine serum albumin
CI: Confidence interval
CO1: Cytochrome c oxidase subunit I
cpds: Complete data set; a data set containing both haploid and diploid individuals before any populations were excluded due to low sample sizes
DNA: Deoxyribonucleic acid
dpds: Diploid data set; a data set containing diploid individuals only
DTT: Dithiothreitol
e.g.: For example
Env only: A model encompassing environmental variables only
FDR: False discovery rate
Fst: Fixation index
Full: A model encompassing geographic distance, resistance distance and environmental variables
GDM: Generalized dissimilarity modelling
Geo only: A model encompassing straight-line geographic distances only
HO: Observed heterozygosity
HPLC: High-performance liquid chromatography
HWE: Hardy-Weinberg equilibrium
i.e.: In other words
IBA: Isolation by adaptation
IBD: Isolation by distance
IBE: Isolation by environment/ecology
IBR: Isolation by resistance
*K*: Number of Structure clusters
k: Number of individuals used for calculating rarefied allelic richnesses
(k)m: (kilo)metre(s)
LAI: Leaf Area Index
LD: (Genotypic) linkage disequilibrium
(μ)l: (Micro)litre
(μ)M: Molar concentration in (micro)mol/litre
MCMC: Marcov chain Monte Carlo
mpds: Mixed-ploidy data set; a data set containing both haploid and diploid individuals
msX: microsatellite loci X
N: Number of individuals
N_A_: Number of alleles
N_E_: Number of effective alleles
PCA: Principal component analysis
PCR: Polymerase chain reaction
PMX: Primer mix X
pop10: Complete data set after excluding populations with fewer than 10 individuals
QSCAT: Quick Scatterometer
r: Pearson’s correlation coefficient
Random: 1,000 random models, each using 16 random environmental variables
Res only: A model encompassing resistance distances only
RGB: Red, green, and blue (colour space)
SDM: Species distribution model
T_m_: Annealing temperature
uH_E_: Unbiased expected heterozygosity
Var: Environmental variable
VIF: Variance inflation factor

## Additional files

### Additional file 1

Table S1: Microsatellite loci used in this work, with original locus names (98), the primer mixes they were assigned to, repeat units (motifs), size ranges, annealing temperatures (Tm), as well as forward and reverse primer sequences (5’–3’).

Table S2: Basic population genetic statistics for each sampling location encompassing diploid (‘dpds’) and both haploid and diploid individuals (‘mpds’). N: number of individuals; N_A_: number of alleles; N_E_: number of effective alleles; A_R_: allelic richness, rarefied for seven individuals (k); uH_E_; unbiased expected heterozygosity; H_O_: observed heterozygosity. Observed heterozygosity values between ‘dpds’ and ‘mpds’ are largely congruent, as SPAGeDi, even though capable of working with mixed-ploidy data sets, does not include haploid individuals in calculating this metric.

Table S3: Multilocus pairwise Fst values derived from ‘dpds’ (below the diagonal) with corresponding FDR-corrected *P* values (above the diagonal). Borderline significant Fst values and *P* values are indicated in bold. See Table S7 for population names.

Table S4: Multilocus pairwise Fst values derived from ‘mpds’ (above the diagonal) with corresponding bootstrap-derived upper 95% confidence intervals below the diagonal. See Table S7 for population names.

Table S5: Environmental variables used, with the ones bolded that were retained after stepwise elimination of variables with a variance inflation factor > 10. Bioclimatic variables (Var 1–19) were obtained from WorldClim v.2 (123, 124), Var 20 (canopy height) from Simard, Pinto (125), vegetation variables (Var 21–25) from The Global Land Cover Facility (126), surface moisture data (Var 26–28) from the BYU Center for Remote Sensing (127), topography variables (Var 29–31) from the Shuttle Radar Topography Mission (128), and human population density (Var 32) from the Socioeconomic Data and Applications Center (129).

Table S6: Pearson correlation coefficients of environmental variables at ‘mpds’ population locations. Values were computed using the ‘cor’ function in R v.3.6.2. See Table S5 for an itemisation of the abbreviations used.

Table S7: Populations sampled, including their abbreviations used in this work, locations, and the number of individuals. ‘cpds’: full data set after exclusion of full siblings and clones; ‘pop10’: ‘cpds’ after excluding populations with fewer than 10 individuals; ‘mpds’: mixed-ploidy data set encompassing both haploid males and diploid females; ‘dpds’: data set containing diploid individuals only.

Table S8: Circuitscape-derived resistance matrix, where higher values indicate higher predicted resistance to inter-population dispersal. See Table S7 for population names.

### Additional file 2

Exemplary Structure output and the GDM splines produced from ‘mpds’ and ‘dpds’.

## Appendix

## References

1. Pritchard JK, Stephens M, Donnelly P. Inference of population structure using multilocus genotype data. Genetics. 2000;155(2):945–59.

2. Guillot G, Mortier F, Estoup A. GENELAND: a computer package for landscape genetics. Mol Ecol Notes. 2005;5(3):712–5.

3. Jombart T, Devillard S, Dufour A-B, Pontier D. Revealing cryptic spatial patterns in genetic variability by a new multivariate method. Heredity. 2008;101(1):92–103.

4. Chen C, Durand E, Forbes F, François O. Bayesian clustering algorithms ascertaining spatial population structure: a new computer program and a comparison study. Mol Ecol Notes. 2007;7(5):747–56.

5. Wright S. Isolation by distance. Genetics. 1943;28(2):114.

6. Adriaensen F, Chardon J, De Blust G, Swinnen E, Villalba S, Gulinck H, et al. The application of ‘least-cost’modelling as a functional landscape model. Landsc Urban Plan. 2003;64(4):233–47.

7. McRae BH. Isolation by resistance. Evolution. 2006;60(8):1551–61.

8. Orsini L, Vanoverbeke J, Swillen I, Mergeay J, De Meester L. Drivers of population genetic differentiation in the wild: isolation by dispersal limitation, isolation by adaptation and isolation by colonization. Mol Ecol. 2013;22(24):5983–99.

9. Forester BR, Jones MR, Joost S, Landguth EL, Lasky JR. Detecting spatial genetic signatures of local adaptation in heterogeneous landscapes. Mol Ecol. 2016;25(1):104–20.

10. Sexton JP, Hangartner SB, Hoffmann AA. Genetic isolation by environment or distance: which pattern of gene flow is most common? Evolution. 2014;68(1):1–15.

11. Hendry AP. Selection against migrants contributes to the rapid evolution of ecologically dependent reproductive isolation. Evol Ecol Res. 2004;6(8):1219–36.

12. Wang IJ, Summers K. Genetic structure is correlated with phenotypic divergence rather than geographic isolation in the highly polymorphic strawberry poison‐dart frog. Mol Ecol. 2010;19(3):447–58.

13. Edelaar P, Alonso D, Lagerveld S, Senar J, Björklund M. Population differentiation and restricted gene flow in Spanish crossbills: not isolation‐by‐distance but isolation‐by‐ecology. J Evol Biol. 2012;25(3):417–30.

14. Nosil P, Egan SP, Funk DJ. Heterogeneous genomic differentiation between walking‐stick ecotypes:“isolation by adaptation” and multiple roles for divergent selection. Evolution: International Journal of Organic Evolution. 2008;62(2):316–36.

15. Hantak MM, Page RB, Converse PE, Anthony CD, Hickerson CAM, Kuchta SR. Do genetic structure and landscape heterogeneity impact color morph frequency in a polymorphic salamander? Ecography. 2019;42(8):1383–94.

16. Myers EA, Xue AT, Gehara M, Cox CL, Davis Rabosky AR, Lemos‐Espinal J, et al. Environmental heterogeneity and not vicariant biogeographic barriers generate community‐wide population structure in desert‐adapted snakes. Mol Ecol. 2019;28(20):4535–48.

17. Manel S, Schwartz MK, Luikart G, Taberlet P. Landscape genetics: combining landscape ecology and population genetics. Trends Ecol Evol. 2003;18(4):189–97.

18. Holderegger R, Wagner HH. A brief guide to Landscape Genetics. Landsc Ecol. 2006;21(6):793–6.

19. Storfer A, Murphy M, Evans J, Goldberg C, Robinson S, Spear S, et al. Putting the ‘landscape’ in landscape genetics. Heredity. 2007;98(3):128.

20. Kraus RH, Van Hooft P, Megens HJ, Tsvey A, Fokin SY, Ydenberg RC, et al. Global lack of flyway structure in a cosmopolitan bird revealed by a genome wide survey of single nucleotide polymorphisms. Mol Ecol. 2013;22(1):41–55.

21. Troast D, Suhling F, Jinguji H, Sahlén G, Ware J. A global population genetic study of *Pantala flavescens*. PLoS ONE. 2016;11(3):e0148949.

22. Geue JC, Thomassen HA. Unraveling the habitat preferences of two closely related bumble bee species in Eastern Europe. Ecology and Evolution. 2020.

23. Hingston AB, Marsden‐Smedley J, Driscoll DA, Corbett S, Fenton J, Anderson R, et al. Extent of invasion of Tasmanian native vegetation by the exotic bumblebee *Bombus terrestris* (Apoidea: Apidae). Austral Ecol. 2002;27(2):162–72.

24. Rasmont P, Coppée A, Michez D, Meulemeester T, editors. An overview of the *Bombus terrestris* (L. 1758) subspecies (Hymenoptera: Apidae). Ann Soc Entomol Fr; 2008: Taylor & Francis.

25. Chapman R, Wang J, Bourke A. Genetic analysis of spatial foraging patterns and resource sharing in bumble bee pollinators. Mol Ecol. 2003;12(10):2801–8.

26. Silva SE, Seabra SG, Carvalheiro LG, Nunes VL, Marabuto E, Mendes R, et al. Population genomics of *Bombus terrestris* reveals high but unstructured genetic diversity in a potential glacial refugium. Biol J Linn Soc. 2020;129(2):259–72.

27. Estoup A, Solignac M, Cornuet J, Goudet J, Scholl A. Genetic differentiation of continental and island populations of *Bombus terrestris* (Hymenoptera: Apidae) in Europe. Mol Ecol. 1996;5(1):19–31.

28. Moreira AS, Horgan FG, Murray TE, Kakouli-Duarte T. Population genetic structure of *Bombus terrestris* in Europe: Isolation and genetic differentiation of Irish and British populations. Mol Ecol. 2015;24(13):3257–68.

29. Lecocq T, Vereecken NJ, Michez D, Dellicour S, Lhomme P, Valterova I, et al. Patterns of genetic and reproductive traits differentiation in mainland vs. Corsican populations of bumblebees. PLoS ONE. 2013;8(6):e65642.

30. Widmer A, Schmid-Hempel P, Estoup A, Scholl A. Population genetic structure and colonization history of *Bombus terrestris* sl (Hymenoptera: Apidae) from the Canary Islands and Madeira. Heredity. 1998;81(5):563.

31. Kraus F, Wolf S, Moritz R. Male flight distance and population substructure in the bumblebee *Bombus terrestris*. J Anim Ecol. 2009;78(1):247–52.

32. Ferrier S, Manion G, Elith J, Richardson K. Using generalized dissimilarity modelling to analyse and predict patterns of beta diversity in regional biodiversity assessment. Divers Distrib. 2007;13(3):252–64.

33. Lecocq T, Rasmont P, Harpke A, Schweiger O. Improving international trade regulation by considering intraspecific variation for invasion risk assessment of commercially traded species: The *Bombus terrestris* case. Conservation Letters. 2016;9(4):281–9.

34. Theodorou P, Radzevičiūtė R, Kahnt B, Soro A, Grosse I, Paxton RJ. Genome-wide single nucleotide polymorphism scan suggests adaptation to urbanization in an important pollinator, the red-tailed bumblebee (*Bombus lapidarius* L.). Proceedings of the Royal Society B: Biological Sciences. 2018;285(1877):20172806.

35. Maebe K, Karise R, Meeus I, Mänd M, Smagghe G. Pattern of population structuring between Belgian and Estonian bumblebees. Scientific reports. 2019;9(1):1–8.

36. Shao Z, Mao H, Fu W, Ono M, Wang D, Bonizzoni M, et al. Genetic Structure of Asian Populations of *Bombus ignitus* (Hymenoptera: Apidae). J Hered. 2004;95(1):46–52.

37. Widmer A, Schmid‐Hempel P. The population genetic structure of a large temperate pollinator species, *Bombus pascuorum* (Scopoli) (Hymenoptera: Apidae). Mol Ecol. 1999;8(3):387–98.

38. Lozier JD, Strange JP, Stewart IJ, Cameron SA. Patterns of range-wide genetic variation in six North American bumble bee (Apidae: *Bombus*) species. Mol Ecol. 2011;20(23):4870–88.

39. Dreier S, Redhead JW, Warren IA, Bourke AF, Heard MS, Jordan WC, et al. Fine‐scale spatial genetic structure of common and declining bumble bees across an agricultural landscape. Mol Ecol. 2014;23(14):3384–95.

40. Koch JB, Looney C, Sheppard WS, Strange JP. Patterns of population genetic structure and diversity across bumble bee communities in the Pacific Northwest. Conserv Genet. 2017;18(3):507–20.

41. Jha S. Contemporary human‐altered landscapes and oceanic barriers reduce bumble bee gene flow. Mol Ecol. 2015;24(5):993–1006.

42. Goulson D, Kaden J, Lepais O, Lye G, Darvill B. Population structure, dispersal and colonization history of the garden bumblebee *Bombus hortorum* in the Western Isles of Scotland. Conserv Genet. 2011;12(4):867–79.

43. Darvill B, Ellis JS, Lye GC, Goulson D. Population structure and inbreeding in a rare and declining bumblebee, *Bombus muscorum* (Hymenoptera: Apidae). Mol Ecol. 2006;15(3):601–11.

44. Amin MR, Than KK, Kwon YJ. Copulation duration of bumblebee *Bombus terrestris* (Hymenoptera: Apidae): Impacts on polyandry and colony parameters. J Asia-Pac Entomol. 2009;12(3):141–4.

45. Zhang H, Zhou Z, Huang J, Yuan X, Ding G, An J. Queen traits and colony size of four bumblebee species of China. Insectes Soc. 2018;65(4):537–47.

46. Duchateau M, Velthuis H. Development and reproductive strategies in *Bombus terrestris* colonies. Behaviour. 1988:186–207.

47. Connop S, Hill T, Steer J, Shaw P. The role of dietary breadth in national bumblebee (*Bombus*) declines: Simple correlation? Biol Conserv. 2010;143(11):2739–46.

48. Wood TJ, Gibbs J, Graham KK, Isaacs R. Narrow pollen diets are associated with declining Midwestern bumble bee species. Ecology. 2019;100(6):e02697.

49. Fontaine C, Collin CL, Dajoz I. Generalist foraging of pollinators: diet expansion at high density. J Ecol. 2008;96(5):1002–10.

50. Avarguès-Weber A, Lachlan R, Chittka L. Bumblebee social learning can lead to suboptimal foraging choices. Anim Behav. 2018;135:209–14.

51. Müller CB, Schmid‐Hempel P. Correlates of reproductive success among field colonies of *Bombus lucorum*: the importance of growth and parasites. Ecol Entomol. 1992;17(4):343–53.

52. Goulson D. Bumblebees: behaviour, ecology, and conservation. Second Edition ed: Oxford University Press; 2009.

53. Knight ME, Martin AP, Bishop S, Osborne JL, Hale RJ, Sanderson RA, et al. An interspecific comparison of foraging range and nest density of four bumblebee (*Bombus*) species. Mol Ecol. 2005;14(6):1811–20.

54. Darvill B, Knight ME, Goulson D. Use of genetic markers to quantify bumblebee foraging range and nest density. Oikos. 2004;107(3):471–8.

55. Ghisbain G, Lozier JD, Rahman SR, Ezray BD, Tian L, Ulmer JM, et al. Substantial genetic divergence and lack of recent gene flow support cryptic speciation in a colour polymorphic bumble bee (*Bombus bifarius*) species complex. Syst Entomol. 2020;45(3):635–52.

56. Alford D. A study of the hibernation of bumblebees (Hymenoptera: Bombidae) in Southern England. The Journal of Animal Ecology. 1969:149–70.

57. Pomeroy N, Plowright R. The relation between worker numbers and the production of males and queens in the bumble bee *Bombus perplexus*. Can J Zool. 1982;60(5):954–7.

58. Müller C, Schmid-Hempel P. Variation in life-history pattern in relation to worker mortality in the bumble-bee, *Bombus lucorum*. Funct Ecol. 1992:48–56.

59. Rundlöf M, Persson AS, Smith HG, Bommarco R. Late-season mass-flowering red clover increases bumble bee queen and male densities. Biol Conserv. 2014;172:138–45.

60. Bowers MA. Resource availability and timing of reproduction in bumble bee colonies (Hymenoptera: Apidae). Environ Entomol. 1986;15(3):750–5.

61. Schmid-Hempel P, Durrer S. Parasites, floral resources and reproduction in natural populations of bumblebees. Oikos. 1991:342–50.

62. Baer B, Morgan ED, Schmid-Hempel P. A nonspecific fatty acid within the bumblebee mating plug prevents females from remating. Proceedings of the National Academy of Sciences. 2001;98(7):3926–8.

63. Duvoisin N, Baer B, Schmid-Hempel P. Sperm transfer and male competition in a bumblebee. Anim Behav. 1999;58(4):743–9.

64. Heinrich B. Bumblebee economics: Harvard University Press; 1979.

65. Vogt FD. Thermoregulation in bumblebee colonies. I. Thermoregulatory versus brood-maintenance behaviors during acute changes in ambient temperature. Physiol Zool. 1986;59(1):55–9.

66. Owen EL, Bale JS, Hayward SA. Can winter-active bumblebees survive the cold? Assessing the cold tolerance of *Bombus terrestris audax* and the effects of pollen feeding. PLoS ONE. 2013;8(11):e80061.

67. Stelzer RJ, Chittka L, Carlton M, Ings TC. Winter active bumblebees (*Bombus terrestris*) achieve high foraging rates in urban Britain. PLoS ONE. 2010;5(3):e9559.

68. Martinet B, Rasmont P, Cederberg B, Evrard D, Ødegaard F, Paukkunen J, et al., editors. Forward to the north: two Euro-Mediterranean bumblebee species now cross the Arctic Circle. Annales de la Société entomologique de France (NS); 2015: Taylor & Francis.

69. Gherghescu D-Ş, Dabija A-M, editors. The Challenges of the Bioclimatic Architecture in Romania. IOP Conference Series: Materials Science and Engineering; 2020: IOP Publishing.

70. Malcheva K, Pophristov V, Marinova T, Trifonova L, editors. Complex approach for classification of winter severity in Bulgaria. AIP Conference Proceedings; 2019: AIP Publishing LLC.

71. Kreyer D, Oed A, Walther-Hellwig K, Frankl R. Are forests potential landscape barriers for foraging bumblebees? Landscape scale experiments with *Bombus terrestris* agg. and *Bombus pascuorum* (Hymenoptera, Apidae). Biol Conserv. 2004;116(1):111–8.

72. Mola JM, Miller MR, O’Rourke SM, Williams NM. Forests do not limit bumble bee foraging movements in a montane meadow complex. Ecol Entomol. 2020;45(5):955–65.

73. Svensson B, Lagerlöf J, Svensson BG. Habitat preferences of nest-seeking bumble bees (Hymenoptera: Apidae) in an agricultural landscape. Agriculture, Ecosystems & Environment. 2000;77(3):247–55.

74. Jha S, Kremen C. Urban land use limits regional bumble bee gene flow. Mol Ecol. 2013;22(9):2483–95.

75. Kadoya T, Washitani I. Predicting the rate of range expansion of an invasive alien bumblebee (*Bombus terrestris*) using a stochastic spatio-temporal model. Biol Conserv. 2010;143(5):1228–35.

76. Liczner AR, Colla SR. A systematic review of the nesting and overwintering habitat of bumble bees globally. J Insect Conserv. 2019;23(5):787–801.

77. Plath O. Notes on the hibernation of several North American bumblebees. Ann Entomol Soc Am. 1927;20(2):181–92.

78. Makinson JC, Woodgate JL, Reynolds A, Capaldi EA, Perry CJ, Chittka L. Harmonic radar tracking reveals random dispersal pattern of bumblebee (*Bombus terrestris*) queens after hibernation. Scientific reports. 2019;9(1):4651.

79. Kämper W, Werner PK, Hilpert A, Westphal C, Blüthgen N, Eltz T, et al. How landscape, pollen intake and pollen quality affect colony growth in *Bombus terrestris*. Landsc Ecol. 2016;31(10):2245–58.

80. Redhead JW, Dreier S, Bourke AF, Heard MS, Jordan WC, Sumner S, et al. Effects of habitat composition and landscape structure on worker foraging distances of five bumble bee species. Ecol Appl. 2016;26(3):726–39.

81. Straub L, Williams GR, Vidondo B, Khongphinitbunjong K, Retschnig G, Schneeberger A, et al. Neonicotinoids and ectoparasitic mites synergistically impact honeybees. Scientific reports. 2019;9(1):1–10.

82. Botías C, Jones JC, Pamminger T, Bartomeus I, Hughes WO, Goulson D. Multiple stressors interact to impair the performance of bumblebee *Bombus terrestris* colonies. J Anim Ecol. 2021;90(2):415–31.

83. Rasmont P, Franzén M, Lecocq T, Harpke A, Roberts SP, Biesmeijer JC, et al. Climatic risk and distribution atlas of European bumblebees: Pensoft Publishers; 2015.

84. Dafni A, Kevan P, Gross CL, Goka K. *Bombus terrestris*, pollinator, invasive and pest: An assessment of problems associated with its widespread introductions for commercial purposes. Appl Entomol Zool. 2010;45(1):101–13.

85. Velthuis HH, Van Doorn A. A century of advances in bumblebee domestication and the economic and environmental aspects of its commercialization for pollination. Apidologie. 2006;37(4):421–51.

86. Semmens T, Turner E, Buttermore R. *Bombus terrestris* (L.) (Hymenoptera: Apidae) now established in Tasmania. J Aust Entomol Soc. 1993;32(4).

87. Matsumura C, Yokoyama J, Washitani I. Invasion status and potential ecological impacts of an invasive alien bumblebee, *Bombus terrestris* L. (Hymenoptera: Apidae) naturalized in Southern Hokkaido, Japan. Glob Environ Res. 2004;8(1):51–66.

88. Macfarlane R, Gurr L. Distribution of bumble bees in New Zealand. N Z Entomol. 1995;18(1):29–36.

89. Romanian Statistical Yearbook. National Institute of Statistics. Bucharest; 2018.

90. Library of Congress.Country profile: Bulgaria. https://www.loc.gov/rr/frd/cs/profiles/Bulgaria.pdf. Accessed 20/05/2019.

91. Milewski P, Szmyd J. Biothermal contrasts while travelling in or between Poland and Bulgaria. EUROPA XXI. 2015;29:73–84.

92. Goulson D, Stout JC. Homing ability of the bumblebee *Bombus terrestris* (Hymenoptera: Apidae). Apidologie. 2001;32(1):105–11.

93. Wolf S, Moritz RF. Foraging distance in *Bombus terrestris* L. (Hymenoptera: Apidae). Apidologie. 2008;39(4):419–27.

94. Smithers CN. The handbook of insect collecting: Their collection, preparation, preservation, and storage: Angus & Robertson; 1988.

95. Campos PF, Gilbert TM. DNA extraction from keratin and chitin. Ancient DNA: Springer; 2012. p. 43–9.

96. Williams PH. Phylogenetic relationships among bumble bees (*Bombus* Latr.): a reappraisal of morphological evidence. Syst Entomol. 1994;19(4):327–44.

97. Wolf S, Rohde M, Moritz RF. The reliability of morphological traits in the differentiation of *Bombus terrestris* and *B. lucorum* (Hymenoptera: Apidae). Apidologie. 2010;41(1):45–53.

98. Stolle E, Rohde M, Vautrin D, Solignac M, Schmid‐Hempel P, Schmid‐Hempel R, et al. Novel microsatellite DNA loci for *Bombus terrestris* (Linnaeus, 1758). Molecular Ecology Resources. 2009;9(5):1345–52.

99. Peakall R, Smouse PE. GenAlEx 6.5: genetic analysis in Excel. Population genetic software for teaching and research—an update. Bioinformatics. 2012;28(19):2537–9.

100. Peakall R, Smouse PE. GENALEX 6: genetic analysis in Excel. Population genetic software for teaching and research. Mol Ecol Notes. 2006;6(1):288–95.

101. Jones OR, Wang J. COLONY: a program for parentage and sibship inference from multilocus genotype data. Molecular ecology resources. 2010;10(3):551–5.

102. Lepais O, Darvill B, O’Connor S, Osborne JL, Sanderson RA, Cussans J, et al. Estimation of bumblebee queen dispersal distances using sibship reconstruction method. Mol Ecol. 2010;19(4):819–31.

103. Bogo G, De Manincor N, Fisogni A, Galloni M, Zavatta L, Bortolotti L. No evidence for an inbreeding avoidance system in the bumble bee *Bombus terrestris*. Apidologie. 2018;49(4):473–83.

104. Kalinowski S. Do polymorphic loci require large sample sizes to estimate genetic distances? Heredity. 2005;94(1):33–6.

105. Van Oosterhout C, Hutchinson WF, Wills DP, Shipley P. MICRO‐CHECKER: software for identifying and correcting genotyping errors in microsatellite data. Mol Ecol Notes. 2004;4(3):535–8.

106. Rousset F. genepop’007: a complete re‐implementation of the genepop software for Windows and Linux. Molecular ecology resources. 2008;8(1):103–6.

107. Hardy O, Vekemans X. SPAGeDi: a versatile computer program to analyse spatial genetic structure at the individual or population levels. Mol Ecol Notes. 2002:618–20.

108. Hurlbert SH. The nonconcept of species diversity: a critique and alternative parameters. Ecology. 1971;52(4):577–86.

109. Kalinowski ST. hp‐rare 1.0: a computer program for performing rarefaction on measures of allelic richness. Mol Ecol Notes. 2005;5(1):187–9.

110. Benjamini Y, Hochberg Y. Controlling the false discovery rate: a practical and powerful approach to multiple testing. Journal of the Royal statistical society: series B (Methodological). 1995;57(1):289–300.

111. R Core Team. R: A language and environment for statistical computing. R Foundation for Statistical Computing. 2019.

112. Clark LV, Jasieniuk M. POLYSAT: an R package for polyploid microsatellite analysis. Molecular Ecology Resources. 2011;11(3):562–6.

113. Falush D, Stephens M, Pritchard JK. Inference of population structure using multilocus genotype data: linked loci and correlated allele frequencies. Genetics. 2003;164(4):1567–87.

114. Puechmaille SJ. The program structure does not reliably recover the correct population structure when sampling is uneven: subsampling and new estimators alleviate the problem. Molecular ecology resources. 2016;16(3):608–27.

115. Pritchard JK, Wen W, Falush D. Documentation for STRUCTURE software: Version 2. 2003.

116. Francis RM. pophelper: an R package and web app to analyse and visualize population structure. Molecular ecology resources. 2017;17(1):27–32.

117. McRae BH, Beier P. Circuit theory predicts gene flow in plant and animal populations. Proceedings of the National Academy of Sciences. 2007;104(50):19885–90.

118. Manion G, Lisk M, Ferrier S, Lugilde KM, Fitzpatrick MC. Package ‘gdm’. A toolkit with functions to fit, plot, and summarize Generalized Dissimilarity Models: CRAN Repository, R. 2017.

119. Thomassen HA, Cheviron ZA, Freedman AH, Harrigan RJ, Wayne RK, Smith TB. Spatial modelling and landscape‐level approaches for visualizing intra‐specific variation. Mol Ecol. 2010;19(17):3532–48.

120. Fitzpatrick MC, Keller SR. Ecological genomics meets community‐level modelling of biodiversity: Mapping the genomic landscape of current and future environmental adaptation. Ecol Lett. 2015;18(1):1–16.

121. Fitzpatrick MC, Sanders NJ, Normand S, Svenning J-C, Ferrier S, Gove AD, et al. Environmental and historical imprints on beta diversity: insights from variation in rates of species turnover along gradients. Proceedings of the Royal Society B: Biological Sciences. 2013;280(1768):20131201.

122. QGIS Development Team. QGIS geographic information system, open source Geospatial Foundation project, version 3.16. 2020.

123. Fick SE, Hijmans RJ. WorldClim 2: new 1‐km spatial resolution climate surfaces for global land areas. International journal of climatology. 2017;37(12):4302–15.

124. WorldClim [Internet] [cited Jan 2018]. Available from: http://www.worldclim.com/version2.

125. Simard M, Pinto N, Fisher JB, Baccini A. Mapping forest canopy height globally with spaceborne lidar. Journal of Geophysical Research: Biogeosciences. 2011;116(G4).

126. The Global Land Cover Facility (GLCF) [Internet] [cited Nov 2014 (LAI/percent tree cover); Nov 2017 (canopy height)]. Available from: Service discontinued.

127. BYU Center for Remote Sensing [Internet] [cited Aug 2015]. Available from: https://www.scp.byu.edu.

128. Shuttle Radar Topography Mission [Internet] [cited Mar 2015]. Available from: https://www2.jpl.nasa.gov/srtm/.

129. Socioeconomic Data and Applications Center (SEDAC) [Internet] [cited May 2015]. Available from: https://sedac.ciesin.columbia.edu/data/collection/gpw-v3.

130. Kelso NV, Patterson T. Introducing natural earth data - naturalearthdata.com. Geographia Technica. 2010;5(82–89):25.

